# A simple method for in situ, multiplexed measurement of RNA degradation by flow cytometry

**DOI:** 10.1101/2021.09.17.460772

**Authors:** Jayan Rammohan, Nina Alperovich, Bin Shao, David Ross

## Abstract

RNA degradation plays a major role in cellular function, but current methods for measuring RNA degradation require RNA purification or are low throughput. Here we show how a flow-FISH assay can be used for high-throughput, *in situ* measurement of RNA degradation without RNA purification. We demonstrate how this approach can be used to simultaneously measure RNA degradation rates of different RNA sequences in a single assay and explore how the assay can be used to examine the effect of cellular context on RNA degradation rates. This assay will be generally useful to quantitatively measure how natural and engineered biological function depends on RNA half-life.

## 1. Introduction

Measuring RNA degradation within cells is critical for mechanistic understanding and engineering of cellular function [1–3]. Unfortunately, RNA degradation techniques that require purification of RNA from cells [4–7], such as RT-qPCR and RNA-seq, introduce uncertainty into these measurements. RNA labeling via fluorescence *in situ* hybridization (FISH) can be used to directly estimate RNA degradation in cells without the need for RNA purification. However, previous uses of FISH for estimating RNA degradation have typically relied on microscopy [8], which limits throughput both in terms of the number of samples analyzed in a given experiment, as well as the number of cells analyzed for each sample.

Here we demonstrate a method for estimating RNA degradation using flow cytometry to detect RNA labeled *in situ* (Fig. 1). This high-throughput approach to measure RNA degradation *in situ* reduces the barrier to RNA degradation measurements by eliminating the needs for RNA purification or low-throughput microscopy. We demonstrate this method using mRNA expressed in *Escherichia coli* (*E. coli*) from the J23101 promoter [9, 10], which is often used as a reference promoter with activity defined as one Relative Promoter Unit (RPU). We show how this approach can be multiplexed to measure degradation of different RNA sequences within a polycistronic transcript. Finally, we demonstrate how this method can be used to measure how RNAseE activity within cells quantitatively affects RNA degradation rates. This work could be expanded to explore the effect of protein-RNA interactions on RNA degradation rates, or to measure RNA degradation of any transcription system in *E. coli* amenable to detection by FISH. We anticipate that this method will be generalizable to other organisms and *in situ* hybridization strategies as well.

**Fig. 1.**
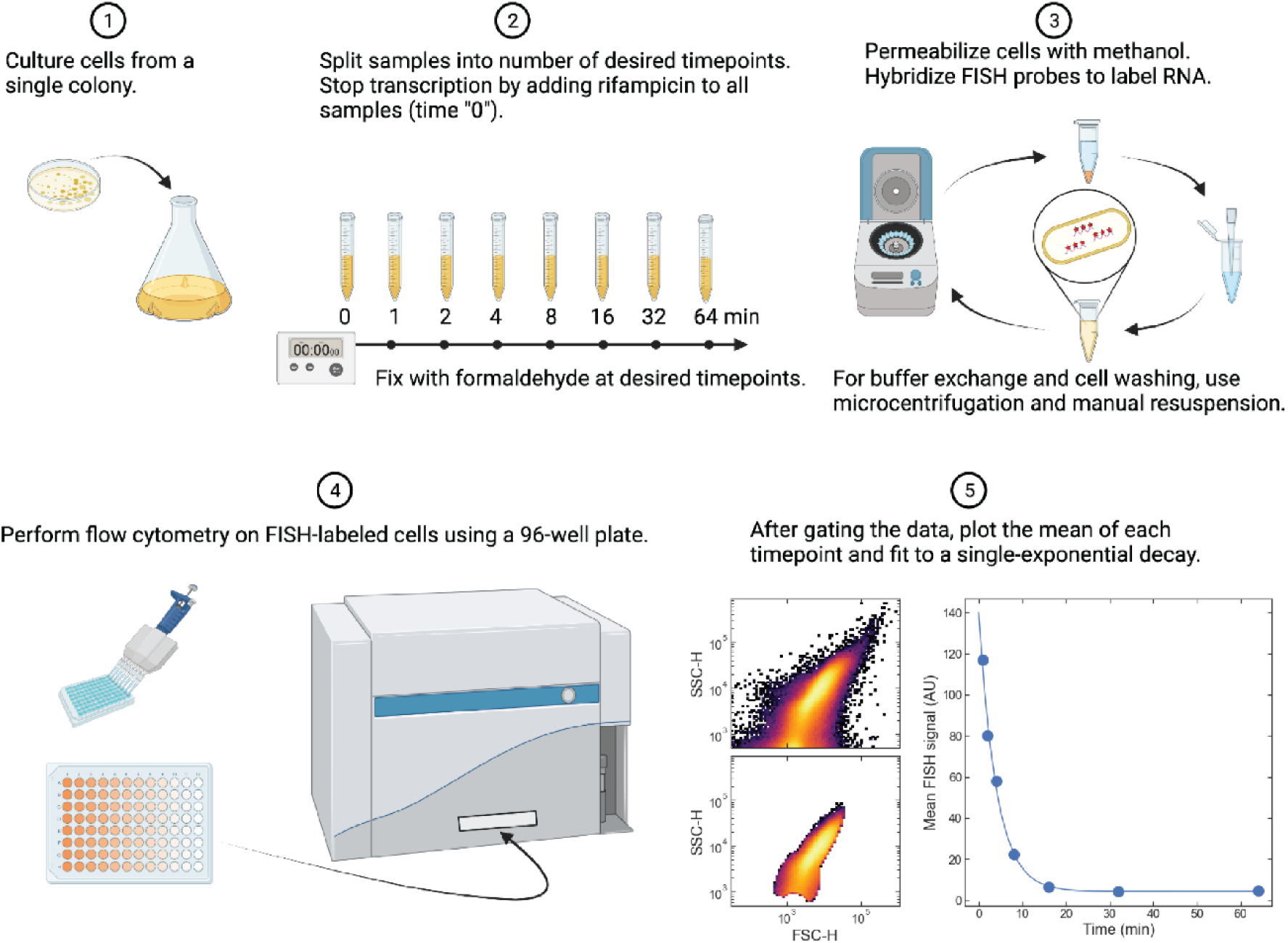
Flow-FISH assay for *in situ* measurement of RNA degradation. (1) To culture cells for estimating RNA degradation by FISH, 80 mL of *E. coli* culture was grown to an OD600 between 0.2-0.3. (2) Cells were split into 8 mL samples and fixed at timepoints following rifampicin addition. (3) Cells were permeabilized and labeled with FISH probes targeting the RNA sequences of interest. (4) Flow cytometry was used to measure the fluorescence per cell at each timepoint. (5) After gating the data (top plot shows raw events, bottom plot shows gated events), the mean FISH signal was plotted versus time (right plot, points), and the data was fit to a single-exponential decay (right plot, line). Example data shown is for FISH-labeled *eyfp* RNA sequence.

## 2. Results

### 2.1 Flow-FISH can be used for *in situ* measurement of RNA degradation

To estimate of RNA degradation rate without RNA purification or microscopy, we developed a flow cytometry assay for measuring FISH-labeled RNA (flow-FISH) in cells following treatment by rifampicin (Fig. 1). RNA degradation in bacteria is commonly estimated by measuring RNA levels at various timepoints following addition of rifampicin to cell culture. Rifampicin stops transcription by stalling bacterial RNA polymerase during transcription initiation, and measurements of RNA levels over time indicate the rate of RNA degradation [4]. In this flow-FISH assay, cells are fixed using formaldehyde at various timepoints following rifampicin addition. Each timepoint then represents a snapshot of the amount of RNA remaining in the cell after transcription has been stopped by rifampicin. Cells are then permeabilized, and FISH probes are added to label a target RNA transcript. Excess FISH probes are removed by washing the cells, and the remaining FISH signal from labeled transcripts within cells are measured by flow cytometry. Mean fluorescence from gated cytometry data decreases over time and can be fit to a single-exponential decay to estimate the RNA degradation rate.

### 2.2 Multiplexed measurement of RNA degradation from different RNA sequences

To demonstrate this assay in a system that is relevant to engineering biology, we used a strain of *E. coli* containing a dual-plasmid system adapted for estimating absolute, quantitative transcriptional flux for engineered genetic circuits in bacteria [10]. This system contains an “expression” plasmid (pSB223 in Fig. 2a), which constitutively expresses a polycistronic transcript with the *eyfp* gene at the 5’ end, and *pp7* hairpin repeats at the 3’ end. The system also contains a “labeling” plasmid (pSB251 in Fig. 2a), which can be induced by anhydrotetracycline (aTc) to express a CFP-PP7 fusion protein that binds the *pp7* repeats in the transcript from the expression plasmid. Although induction of the labeling plasmid was not performed in this study, those experiments are an obvious extension of this work, and a comparison between the induced versus non-induced labeling system would provide insight into the effect of protein-RNA interactions on RNA degradation rate. As a negative control, we also tested a strain of *E. coli* containing a plasmid that does not express fluorescent protein.

**Fig. 2.**
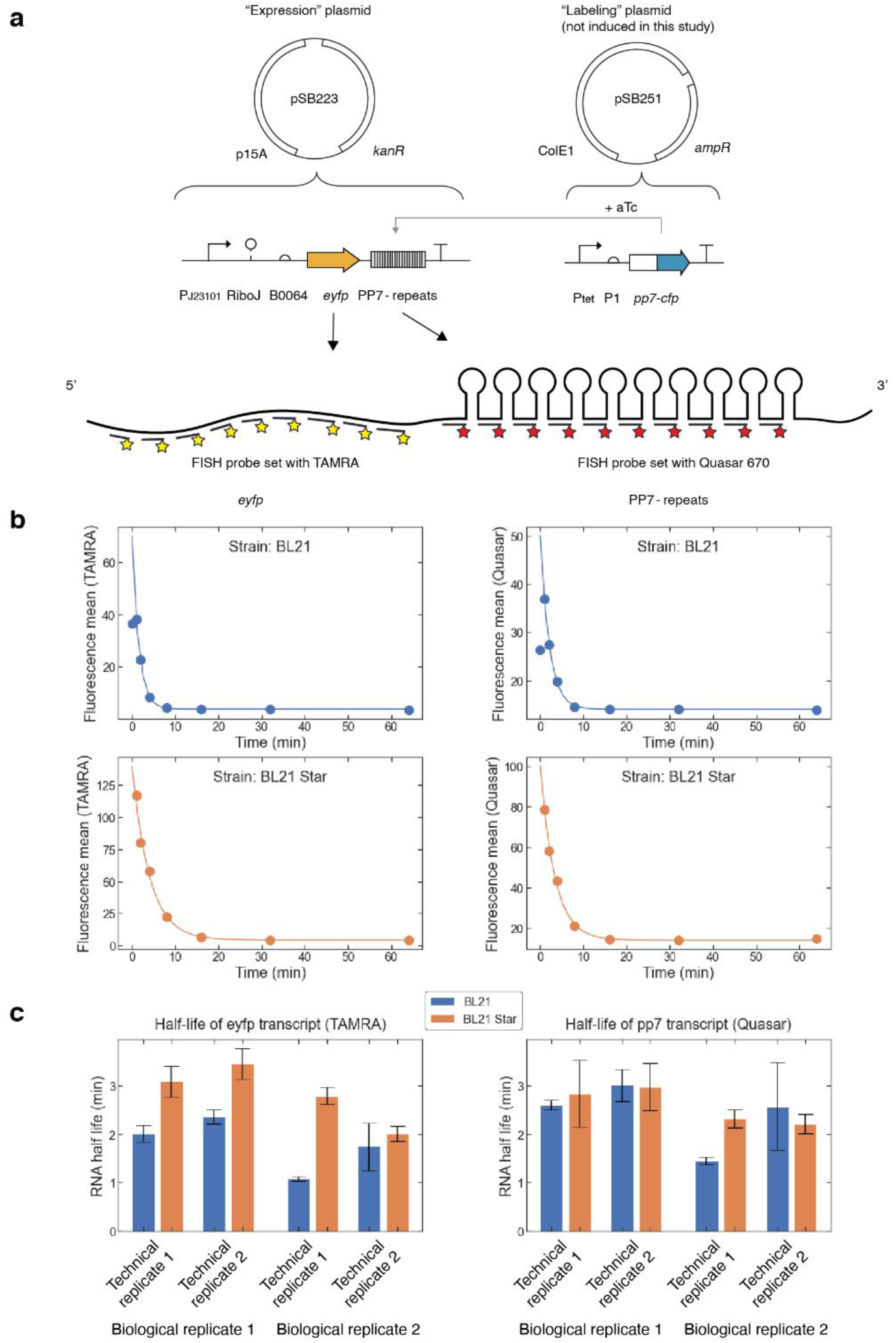
Multiplexed flow-FISH can be used to measure the effect of RNAseE activity on RNA degradation rate of multiple target RNA sequences. (a) A polycistronic transcript expressed as part of a dual-plasmid expression was targeted by two different sets of FISH probes. The “labeling” plasmid was not induced by aTc in this study. (b) RNA decay curves for *eyfp* and *pp7* sequences in BL21 and BL21 Star cells. Points represent mean of single-cell fluorescence data. Solid lines represent fits to single-exponential decay starting from 1 min. (c) Half-life of e*yfp* sequence (left) and *pp7* sequence (right) in BL21 (blue) and BL21 Star (orange) cells. Technical replicates were performed using flow cytometry plates prepared on different days using the original sample preparation. Two biological replicates were performed on different days. Error bars represent one standard deviation based on the non-linear least-squares fits.

To detect RNA degradation rates from different sequences within a polycistronic transcript, we cultured the pSB223 and pSB251 strains as described in the methods. Following rifampicin treatment, we fixed cells at various timepoints (0 min, 1 min, 2 min, 4 min, 8 min, 16 min, 32 min, and 64 min). For all samples, we targeted the *eyfp* sequence with a set of TAMRA-labeled FISH probes, and we targeted the *pp7* repeat sequence with a set of Quasar-670-labeled FISH probes (Fig. 2b). For each sequence, we detected a change in fluorescence over time following rifampicin addition. Because transcript levels can initially rise following rifampicin addition [7], we used timepoints starting from 1 min onward to fit a single-exponential decay. Using two different strains (BL21 and BL21 Star, see below), we performed two biological replicates, each started from a single colony. For each sample, we performed two technical replicates (flow cytometry measurements run on different days using the same labeled samples). For all technical and biological replicates, we observed a decrease in FISH signal with time, for both the *eyfp* and *pp7* sequences (Figs. 2–6).

In the BL21 strain, the half-life of the *pp7* sequence was consistently longer than the half-life of the *eyfp* sequence (Fig. 2c, see also Figs. 3–6). In the BL21 Star strain, on the other hand, the *pp7* and *eyfp* sequences had similar decay rates (within the measurement uncertainty). These results suggest that the two RNA sequences (*eyfp* and *pp7*) can degrade at different rates, despite being part of the same RNA transcript.

**Fig. 3.**
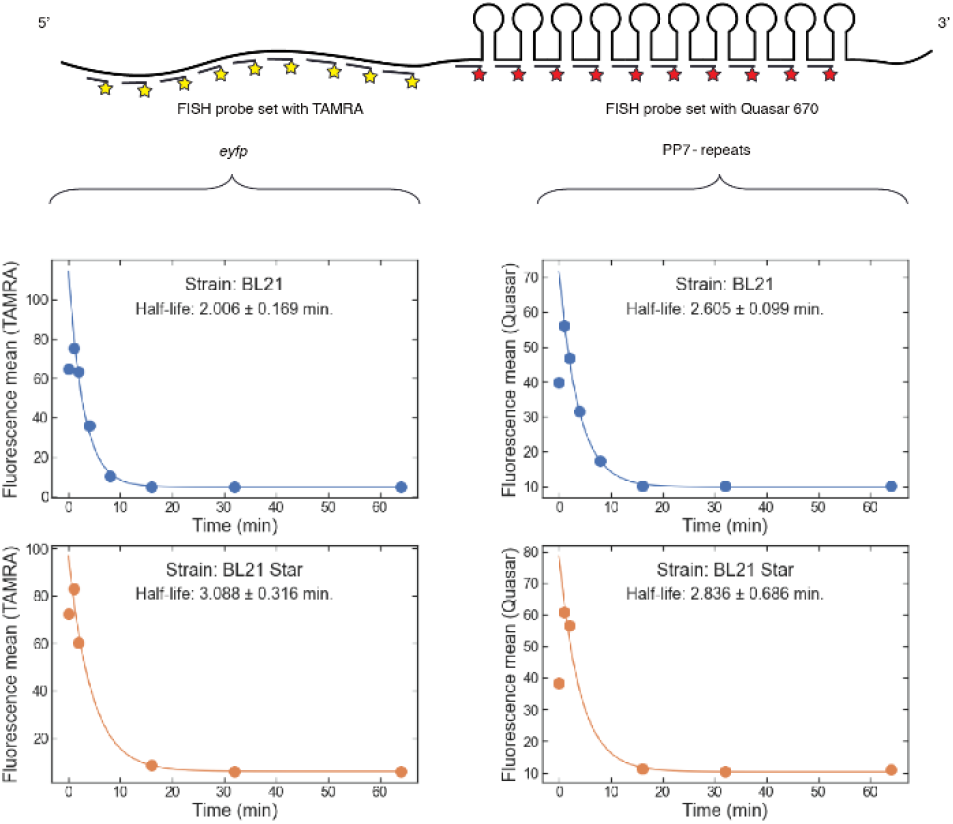
Flow-FISH RNA decay curves for biological replicate 1, technical replicate 1. For all plots, points represent fluorescence mean for samples at each timepoint. Solid lines represent single-exponential fits starting from 1 min. Plots on left are measurements from TAMRA-labeled FISH probes targeted to the *eyfp* sequence. Plots on right are measurements from Quasar 670 FISH probes targeted to the *pp7* repeat sequence. For each FISH probe set, data is shown for the BL21 strain (top, data shown in blue) or BL21 Star strain (bottom, data shown in orange). Based on the fit, the estimated half-life and one-standard-deviation uncertainty are shown in each plot.

**Fig. 4.**
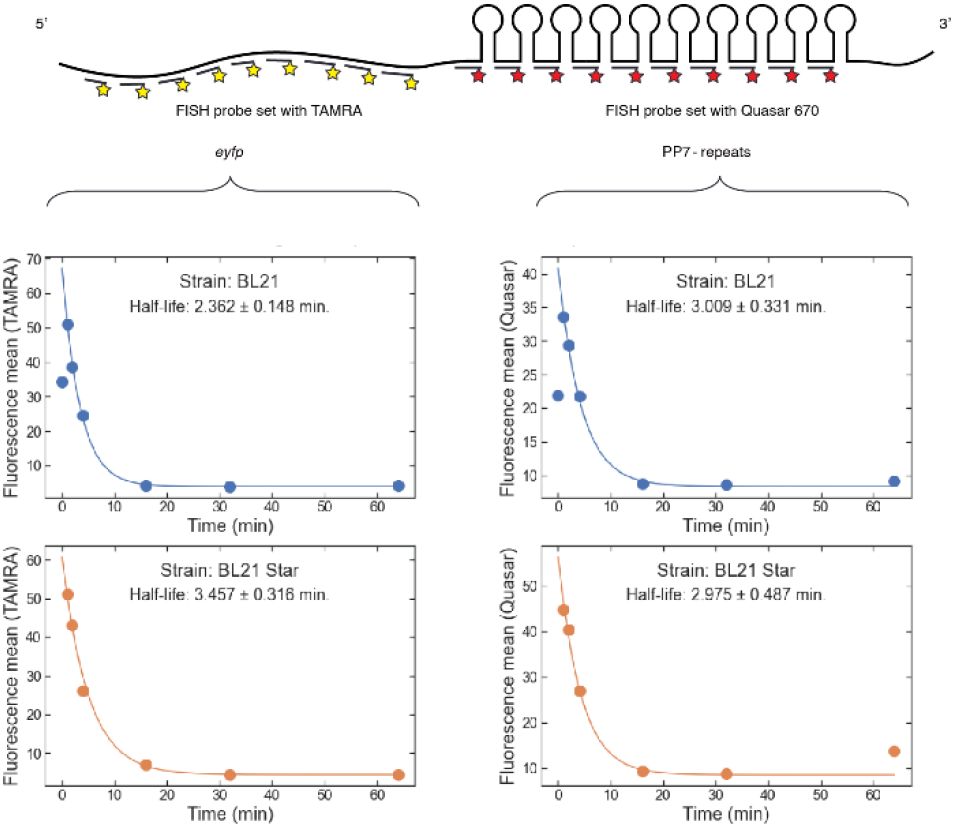
Flow-FISH RNA decay curves for biological replicate 1, technical replicate 2. For all plots, points represent fluorescence mean for samples at each timepoint. Solid lines represent single-exponential fits starting from 1 min. Plots on left are measurements from TAMRA-labeled FISH probes targeted to the *eyfp* sequence. Plots on right are measurements from Quasar 670 FISH probes targeted to the *pp7* repeat sequence. For each FISH probe set, data is shown for the BL21 strain (top, data shown in blue) or BL21 Star strain (bottom, data shown in orange). Based on the fit, the estimated half-life and one-standard-deviation uncertainty are shown in each plot.

**Fig. 5.**
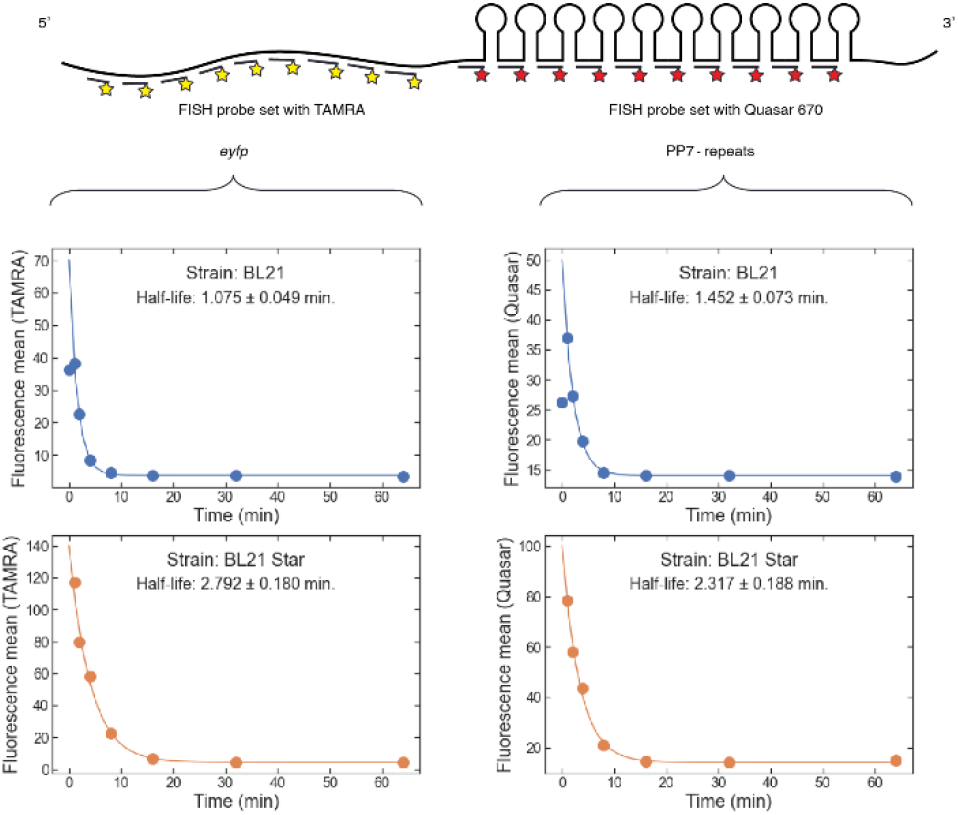
Flow-FISH RNA decay curves for biological replicate 2, technical replicate 1. For all plots, points represent fluorescence mean for samples at each timepoint. Solid lines represent single-exponential fits starting from 1 min. Plots on left are measurements from TAMRA-labeled FISH probes targeted to the *eyfp* sequence. Plots on right are measurements from Quasar 670 FISH probes targeted to the *pp7* repeat sequence. For each FISH probe set, data is shown for the BL21 strain (top, data shown in blue) or BL21 Star strain (bottom, data shown in orange). Based on the fit, the estimated half-life and one-standard-deviation uncertainty are shown in each plot.

**Fig. 6.**
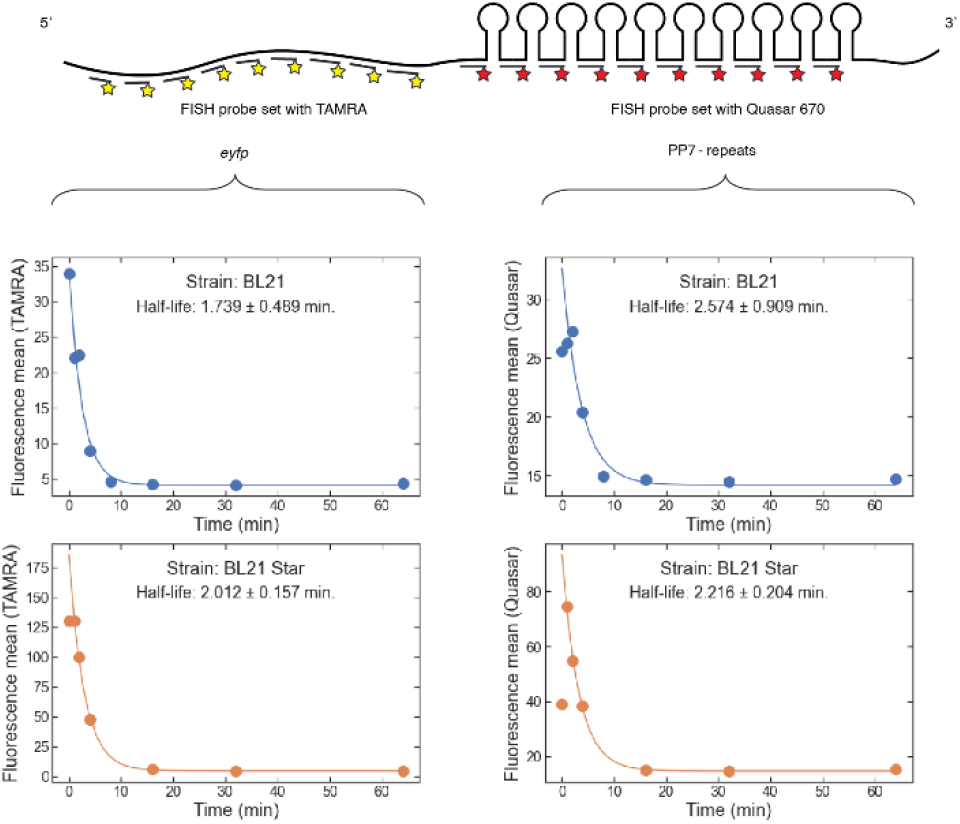
Flow-FISH RNA decay curves for biological replicate 2, technical replicate 2. For all plots, points represent fluorescence mean for samples at each timepoint. Solid lines represent single-exponential fits starting from 1 min. Plots on left are measurements from TAMRA-labeled FISH probes targeted to the *eyfp* sequence. Plots on right are measurements from Quasar 670 FISH probes targeted to the *pp7* repeat sequence. For each FISH probe set, data is shown for the BL21 strain (top, data shown in blue) or BL21 Star strain (bottom, data shown in orange). Based on the fit, the estimated half-life and one-standard-deviation uncertainty are shown in each plot.

### 2.3 Effect of RNAseE activity on RNA degradation rates

RNAseE is responsible for multiple RNA degradation pathways within the cell, and *E. coli* strains containing RNAseE variants are commonly used to increase protein yields. For example, the BL21 strain of *E. coli* is offered as a “Star” version which contains the rne131 RNAseE variant without a C-terminal domain. The RNAseE C-terminal domain acts as a scaffold for assembly of the degradasome complex. The absence of the C-terminal domain is thought to generally increase RNA lifetimes within the cell and can result in increased expression of proteins such as beta-galactosidase.

To examine the effect of RNAseE activity on RNA degradation rate, we compared RNA degradation rates between BL21 and BL21 Star strains. For the *eyfp* sequence, the RNA degradation rate in the BL21 Star strain was consistently slower than in the BL21 strain (Fig. 2c). For the *pp7* stem loop RNA sequence, however, the degradation rates in BL21 and BL21 Star were within the measurement uncertainty. These results suggest that changes in RNAseE activity can have different impacts on the degradation of different regions of a transcript.

## 3. Discussion

We have demonstrated a simple flow-FISH assay for measuring RNA degradation rate *in situ*, circumventing the need for RNA purification or microscopy. Using different FISH probe sets, this assay can be multiplexed for estimating RNA degradation of different RNA sequences in a single sample preparation. The data shown here suggest that this assay would potentially be useful for investigating the effects of RNAseE activity on RNA degradation rate. Such direct, quantitative measurements of RNA degradation represent progress towards mechanistic understanding and engineering of bacteria.

While we have demonstrated the flow-FISH assay in *E. coli*, this protocol could be adapted to estimate RNA degradation in other organisms. For example, the type 3 secretion system in Salmonella is influenced by RNA stability [11]. In addition, the flow-FISH assay could be used to measure the effects of RNA sequence on engineered function for biomanufacturing applications. These quantitative measurements could be useful for characterizing new tools for engineering biology, such as degradation-tuning RNAs [3]. Finally, the assay could be easily adapted for FISH variants such as hybridization chain reaction (HCR) [12], which could potentially offer a way to improve the assay’s signal-to noise-ratio using long amplification times. Taken together, this method represents a straightforward and versatile tool for measuring RNA degradation in bacteria.

## 4. Methods

### 4.1 RNA degradation assay and FISH sample preparation

*E. coli* strains BL21 (DE3) (Invitrogen) or BL21 Star (DE3) (Invitrogen), transformed with plasmids pSB223 and pSB251, were streaked on LB agar plates (Miller, BD Difco) with antibiotics (34 µg/mL kanamycin and 100 µg/mL ampicillin) and grown overnight at 37 °C. For cell culture, we used a rich variant of M9-glycerol media containing M9 salt (Fisher Scientific), 4 % ultrapure glycerol (Sigma-Aldrich), 2 % casamino acids (Millipore Sigma), 2 mmol/L MgSO4 (Millipore Sigma), 0.1 mmol/L CaCl2 (Millipore Sigma), and 0.034 g/L thiamine hydrochloride (Sigma-Aldrich). Single colonies were inoculated into 3 mL of rich M9- glycerol media with kanamycin (34 µg/mL, Gold Biotechnology) and ampicillin (100 µg/mL, Gold Biotechnology) in 14 mL round-bottom culture tubes. The cells were grown overnight for 16 h at 37 °C in an incubator shaking at 300 rpm (Benchmark Scientific). Then, the cultures were diluted 200-fold into two 40 mL cultures, each in 250 mL conical flasks. The cultures were incubated at 37 °C for approximately 4 h shaking at 300 rpm. When the measured optical density at 600 nm (OD600) was between 0.2 and 0.25, 8 mL of each culture was aliquoted into 10 different 15 mL centrifuge tubes (Falcon) kept at 37 °C shaking at 300 rpm (Eppendorf ThermoMixer C with 15 mL SmartBlock). For each of the 8 mL samples, 160 μL of 25 mg/mL rifampicin (Sigma-Aldrich) was added and vortexed for 3 s to mix, and a separate timer was started for each individual sample to track the time after rifampicin addition. To fix the cells and stop RNA degradation at different time points following rifampicin addition, 4 mL of a formaldehyde solution (3.7% by weight, in 1x phosphate buffered saline, PBS) was added to each culture and mixed by pipetting. After cultures for all time points were fixed for 30 min at room temperature, cells were washed twice with 1x PBS, resuspended in 85% methanol for permeabilization for 1 h at room temperature, and then stored at 4 °C for 16 hours. Cells were transferred to new microcentrifuge tubes, washed in a solution of 50 % formamide Wash Buffer A (Biosearch Technologies), and then each of the 21 samples were resuspended in 50 μL of 50 % formamide Hybridization Buffer (Biosearch Technologies) containing the appropriate FISH probe mix. Hybridized samples were incubated at 30 °C for 16 h to label RNA. Cells were washed 3 times with 50 % formamide Wash Buffer A and incubated at 30 °C for 30 min between each wash. Cells were then washed in 500 μL Wash Buffer B (Biosearch Technologies), and resuspended in 50 μL freshly filtered 5× saline-sodium citrate (SSC) buffer (Ambion) for flow cytometry.

Additional details are provided in Appendix A (growth of cultures), Appendix B (step by step protocol), and Appendix C (sequences of FISH probe sets).

### 4.2 Flow cytometry and analysis

Samples were pipetted into a 96 well U-bottom plate at approximately 1:100 dilution by adding 2 μL of sample into 198 μL freshly filtered 5x SSC. Cytometry was performed using a CytoFLEX flow cytometer (Beckman Coulter) with a 561 nm excitation laser and 585/42 nm bandpass filter for TAMRA detection, and a 638 nm excitation laser and 660/10 nm bandpass filter for Quasar 670 detection. Analysis was performed using an automated gating algorithm [13] to distinguish singlet cell events from background and multi-plet cell events. For each replicate, the mean TAMRA and Quasar 670 signals at each time point were fit to a single exponential decay to estimate the half-lives for each RNA sequence. Weighted non-linear least squares fitting was used, with the residuals weighted by the inverse of the squared standard error of the mean (SEM) for each signal. Some samples had an unusually low number of detected singlet cell events (typically 2-4 samples per replicate); samples with fewer than 10,000 detected singlet events were excluded from the analysis.

### 4.3 Data availability

The data for all replicates is available through the NIST Data Portal: https://data.nist.gov/od/id/mds2-2437, doi:10.18434/mds2-2437.

## 5. Appendix A: Growth protocol

Presented as recommended by the Minimum Information Standard for Engineering Organism Experiments (MIEO) [14].

Media Components
| Component | Concentration<br>(prepared in DI<br>water, 18 MΩ-cm) |
| --- | --- |
| Potassium phosphate | 3 g/L |
| Disodium phosphate | 6.78 g/L |
| Sodium chloride | 0.5 g/L |
| Ammonium chloride | 1.0 g/L |
| Glycerol | 40 g/L |
| Casamino acids | 20 g/L |
| Calcium chloride | 0.0147 g/L |
| Magnesium sulfate | 4.93 g/L |
| Vitamin B1 (Thiamine) | 0.034 g/L |

Container Geometry
| *Overnight growth* |  |
| --- | --- |
| Container Type | Culture tube |
| Container shape | Round |
| Container bottom | Round |
| Container volume | 14 mL |
| Fill volume | 14 % (2 mL) |
| Cover | Snap cap (plastic) |
| *Outgrowth* |  |
| Container Type | Shake flask |
| Container shape | Conical |
| Container bottom | Baffled |
| Container volume | 250 mL |
| Fill volume | 16 % (40 mL) |
| Cover | Snap cap (metal) |

Container Shaking
| *Overnight growth* |  |
| --- | --- |
| Shaking speed | 300 rpm |
| Shaking diameter | 2.5 cm |
| Shaking mode | Orbital |
| *Outgrowth* |  |
| Shaking speed | 300 rpm |
| Shaking diameter | 2.5 cm |
| Shaking mode | Orbital |

Time
| *Overnight growth* |  |
| --- | --- |
| Growth time | 16 hours |
| *Outgrowth* |  |
| Growth time | 4 hours |

Environment
| *Overnight growth* |  |
| --- | --- |
| Temperature | 37 °C |
| *Outgrowth* |  |
| Temperature | 37 °C |

Inoculum
| *Overnight growth* |  |
| --- | --- |
| Type | Single colony |
| Concentration at inoculation | N/A |
| *Outgrowth* |  |
| Type | Culture |
| Concentration at inoculation | Optical density at 600 nm<br>= $0.24 \pm 0.01$ * |

\*Average ± standard deviation, all samples and replicates.

Selective agents
| *Overnight growth* |  |
| --- | --- |
| Antibiotic | Concentration |
| Kanamycin (Kan) | 34 µg/mL |
| Ampicillin (Amp) | 100 µg/mL |
| *Outgrowth* |  |
| Antibiotic | Concentration |
| Kanamycin (Kan) | 34 µg/mL |
| Ampicillin (Amp) | 100 µg/mL |

## 6. Appendix B: Step-by-step protocol for RNA degradation estimated by flow-FISH

### 6.1 Growth protocol for cell culture

1. Day 1 – streak plate(s)
2. Day 2 – pick individual colonies for overnight growth in 2 mL cultures, shaking and incubating at 37 °C
3. Day 3 – dilute overnight cultures 1:200 (400 μL into 80 mL), divide the culture equally between two 250 mL baffled culture flasks, and incubate at 37 °C (with shaking) until the OD_600_ is between 0.2 and 0.25. This usually takes between 3 hours and 4 hours.

### 6.2 RNA degradation sample preparation

1. Label 15 mL screw top culture tubes with each timepoint to be measured.
2. Set up Eppendorf ThermoMixer C with 15 mL SmartBlock and set the temperature to 37 °C. Place empty, labeled 15 mL screw top culture tubes in the ThermoMixer.
3. When the cultures in the culture flasks reach an OD_600_ of approximately 0.2, remove from incubator, and for each of the timepoints, aliquot 8 mL into a 15 mL tube and place tubes in the ThermoMixer with 15 mL SmartBlock shaking at 300 rpm.
4. For the zero timepoint, add 4 mL formaldehyde solution (3.7 % in 1x PBS) premixed with 160 μL of rifampicin into the appropriate 8 mL culture and pipette up and down several times to mix (final formaldehyde concentration is approximately 1.2 %). This can be done using a 5 mL capacity pipette.
5. For all other timepoints, add 160 μL of 25 mg/mL rifampicin into the appropriate 8 mL culture, vortex to mix, start a countdown timer for the appropriate time (in this case, 1 min, 2 min, 4 min, 8 min, 16 min, 32 min, or 64 min), and place the culture back in the ThermoMixer.
  a. Plan on approximately 30 sec for each rifampicin addition (add rifampicin, vortex mix, and start countdown timer).
  b. It is useful to start by adding rifampicin to the longest timepoints first, and work backwards (in this case, first add rifampicin to the 64 min timepoint, then the 32 min timepoint, and so on).
  c. As each countdown timer expires, add 4 mL formaldehyde solution (3.7 % working stock) to the 8 mL culture, and mix by pipetting and inverting tube.
  d. Allow all samples at least 30 min at room temperature after formaldehyde addition.

### 6.3 FISH sample preparation

1. Starting with fixed cells, centrifuge (4000 g for 10 min) at room temperature, remove supernatant and resuspend in 200 μL 1x PBS.
2. Transfer 200 μL of sample into 1.7 mL microcentrifuge tubes, add 800 μL 1x PBS.
3. Centrifuge cultures at 1500 g for 7 min.
4. Remove supernatant and gently resuspend cell pellet in 200 μL of 1x PBS, then add 800 μL.
5. Centrifuge cultures at 1500 g for 7 min.
6. Remove supernatant and gently resuspend cell pellet in 150 μL of 1x PBS, then add 850 μL 100% methanol for permeabilization.
7. Incubate samples at room temperature for 1 hour. Then store samples at 4 °C overnight.
8. Transfer all samples to new microcentrifuge tubes. Spin at 600 g for 7 min then remove supernatant (85% methanol). Gently resuspend each cell pellet in Wash Buffer A (with 50% formamide), first using 200 μL, then add 800 μL.
9. Label RNA (this step takes between 2 hours and 3 hours, depending on the number of samples)
10. Move samples to new 1.7 mL microcentrifuge tubes.
11. Centrifuge samples at 600 g for 7 min. Then, remove supernatant and resuspend in Wash Buffer A (with 50 % formamide). For resuspending pellet, first use 100 μL, then add an additional 900 μL.
  a. Centrifuge samples at 1500 g for 7 min. Then, remove supernatant and resuspend in 50 μL Probe Mix (1 μmol/L in Hybridization Buffer with 50 % formamide). For multiplex detection, FISH probe sets can be added together at this step.
  b. Incubate in the dark at 30 °C overnight (approximately 16 hours).
12. Wash samples 3 times in Wash Buffer A (with 50 % formamide) and incubate at 30 °C for 30 min between each wash.
13. Add 400 μL Wash Buffer A to each 50 μL sample in Probe Mix.
14. Centrifuge samples at 600 g for 3.5 min. Remove supernatant – pellet is easy to lose at this point.
15. Resuspend pellet in in two steps, first adding 50 μL and then 150 μL Wash Buffer A. Using 50 μL provides better control over the initial pellet resuspension, then add additional 150 μL, and mix by pipetting to wash.
16. Incubate in the dark at 30 °C for 30 min.
17. Repeat steps b-d.
18. Wash cells before measurement
19. Centrifuge samples at 600 g for 3.5 min, remove supernatant, resuspend in in two steps, first adding 50 μL and then 450 μL Wash Buffer B.
20. Centrifuge samples at 600 g for 3.5 min, remove supernatant, resuspend in 50 μL freshly filtered 5x SSC.

### 6.4 Flow cytometry and analysis

1. Prepare a 96-well plate for cytometry measurement. Ideally, before preparing flow cytometry plate, use a diluted tube measurement to estimate cell concentration from each timepoint, and dilute appropriately to achieve approximately 10^5^ events per timepoint. Based on yields, 1:100 dilution typically provides a sufficient number of cells for analysis.
2. Optimize gain for each fluorescence channel using sample with highest expected fluorescence.
3. To avoid (and measure) carryover between samples, include at least 2 wells of freshly filtered buffer in between each sample well.
4. Export data to Flow Cytometry Standard (FCS) format for analysis.
5. Analyze cytometry data. Use semi-automated gating [13] and estimate mean fluorescence from all samples.
6. Estimate RNA degradation rate. For each gated distribution, plot mean FISH signal versus time. Assuming RNA decay can be estimated using a single-exponential decay, fit the points to the formula 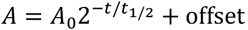.
  a. Because transcript levels can rise immediately following rifampicin addition, use only data for time ≥ 1 min for fit.

## 7. Appendix C: Sequences of FISH Probe Sets

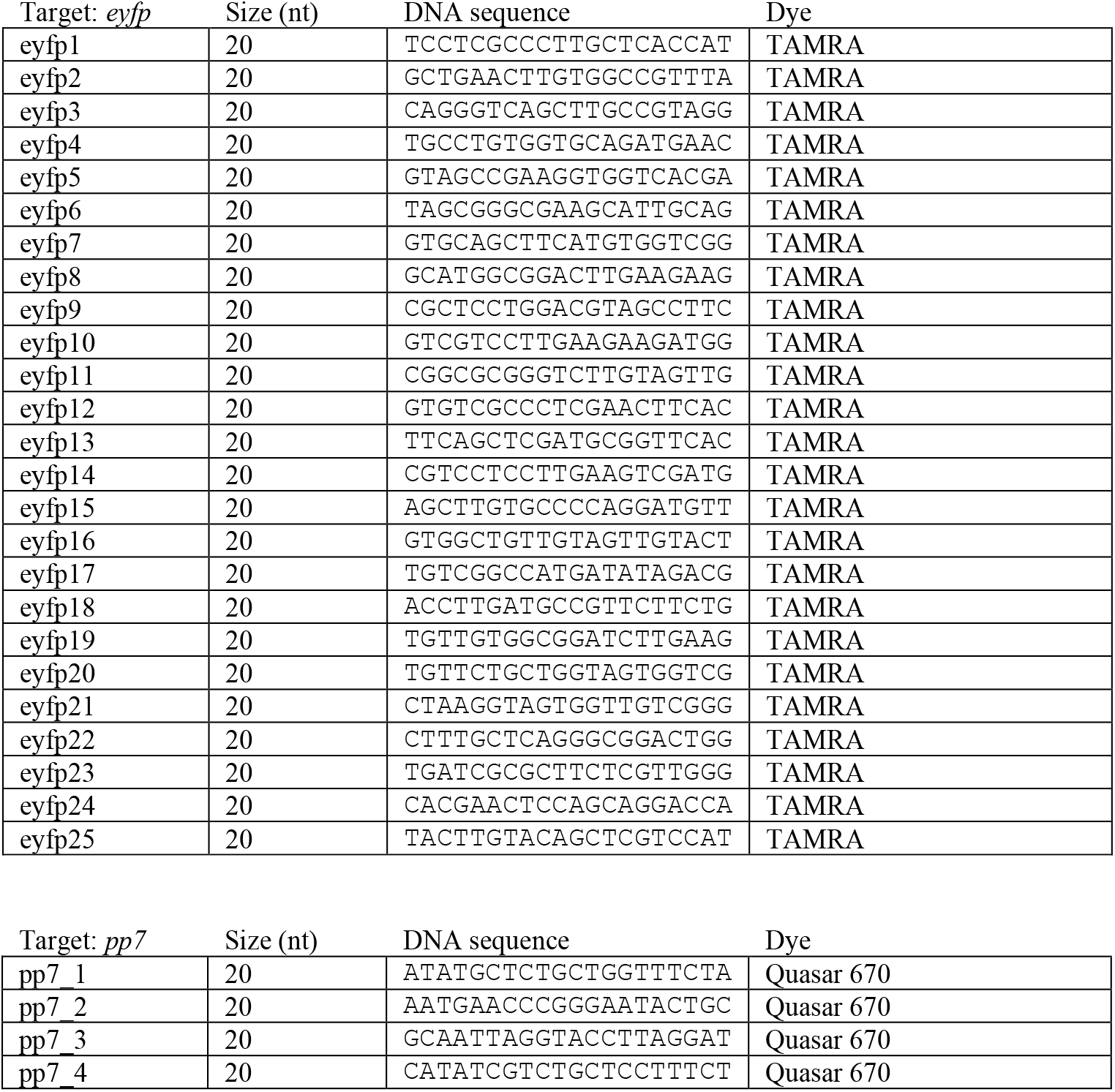

## Acknowledgments

The authors thank Christina Bergonzo, Danielle Tullman-Ercek, Julie Liang, Kirsten Parrat, Eugenia Romantseva, Samuel Schaffter, and Elizabeth Strychalski for useful discussions regarding this work.

## About the authors

*Jayan Rammohan is a research chemist in the Cellular Engineering Group of the Biosystems and Biomaterials Division of the Materials Measurement Laboratory at NIST. His research focuses on transcription metrology to meet the needs of the engineering biology community*.

*Nina Alperovich is a biological science technician in the Cellular Engineering Group of the Biosystems and Biomaterials Division of the Materials Measurement Laboratory at NIST. She is a chemical engineer and developed her career as a researcher in the fields of biochemistry, microbiology and engineering biology*.

*Bin Shao is a postdoctoral fellow at the Department of Molecular and Cellular Biology at Harvard University. His research focuses on the development of single-cell level quantitative methods for engineering biology*.

*David Ross is a physicist in the Cellular Engineering Group of the Biosystems and Biomaterials Division of the Materials Measurement Laboratory at NIST. He uses large-scale measurements and laboratory evolution to develop methods for predictive engineering of biological function*.

*The National Institute of Standards and Technology is an agency of the U.S. Department of Commerce*.

